# Recombinant *Echis* disintegrins reveal distinct inhibitory profiles on platelet aggregation and endothelial cell migration

**DOI:** 10.64898/2026.09.05.749579

**Authors:** Iara Aimê Cardoso, Sophie Hall, Thomas Crasset, Mark C. Wilkinson, Adam Robinson, Bronwyn Rand, Dakang Shen, Connor Webb, Richard Stenner, David Waterhouse, Ingeborg Hers, Renaud Vincentelli, Imre Berger, Nicholas R. Casewell, Loïc Quinton, Alastair W. Poole, Christiane Schaffitzel

## Abstract

Disintegrins are snake venom toxins that bind integrins and modulate platelet aggregation and cell migration. Isolation from venom often yields insufficient quantities for comprehensive study, making recombinant expression essential for detailed toxin characterisation. Here, we produced and characterised four disintegrins from *Echis coloratus* and *Echis ocellatus* in *Escherichia coli*. These homologous proteins contain distinct integrin-binding motifs: EcDis_RGD, EoDis_RGD, EcDis_KGD, and EcDis_VGD. Functional activities were evaluated using human platelet aggregation and endothelial cell scratch wound-healing assays. RGD-containing disintegrins exhibited the strongest inhibition of platelet aggregation, with EcDis_RGD showing the highest potency. The EcDis_KGD disintegrin inhibited platelet aggregation less potently, whereas EcDis_VGD showed no detectable activity in our assays. In endothelial cells, EcDis_RGD disintegrin markedly impaired cell migration, while EoDis_RGD displayed weaker anti-migratory activity. Neither KGD-nor VGD-containing disintegrins significantly affected wound closure. EoDis_RGD exhibited similar antiplatelet activity to the native, venom-purified disintegrin ocellatusin, validating our recombinant expression strategy. However, the corresponding PII-SVMP preparation displayed reduced inhibitory activity, consistent with incomplete generation of fully mature disintegrins species. Our findings demonstrate the importance of both integrin-binding motif identity and surrounding sequence context in determining disintegrin function, and support further use of recombinant toxins in toxinology and as therapeutic lead molecules.

## 1. Introduction

Snake venom is a highly complex mixture of biologically active toxins, which produce a broad spectrum of clinical effects, including both local and systemic manifestations in snakebite patients (Gutiérrez et al., 2017). Despite its toxicity, snake venom is increasingly recognised as a valuable source of therapeutic agents, with each venom containing hundreds of pharmacologically active components capable of targeting diverse physiological systems in the human body (Mohamed Abd El-Aziz et al., 2019).

Disintegrins are a family of small, cysteine-rich, non-enzymatic venom toxins ranging in size from 4 to 15 kDa. These proteins originate either from the proteolytic processing of snake venom metalloproteinases (SVMPs) or via direct translation from mRNAs that lack the metalloprotease-coding region (also known as ‘short-coding disintegrins’ or ‘true disintegrins’) (Juárez et al., 2006; Okuda et al., 2002). Snake venom disintegrins interact specifically with integrins, a group of cell adhesion receptors on the cell surface, including platelet, vascular endothelial, and certain tumour cells (Calvete et al., 2005; Cesar et al., 2019; Kini & Evans, 1992). During envenoming, they exhibit a wide array of functions, notably binding to platelet receptors, which inhibits their aggregation and results in disruption of the haemostatic process (Almeida *et al*., 2023). Integrins are critical mediators of cell adhesion, wound repair, and neuronal connectivity, while also contributing to a range of pathological conditions including inflammation, fibrosis, atherosclerosis, and cancer progression (Cesar *et al*., 2019). The structural determinant of selective integrin binding is a short tripeptide motif, such as RGD (arginine-glycine-aspartate), typically located within a 9 to 11 residue loop projecting from the protein core. Although RGD is most common, other motifs such as KGD, MGD, VGD, WGD, MLD, KTS, and RTS have also been identified (Almeida et al., 2023; Calvete et al., 2005; Ruoslahti, 1996; Vasconcelos et al., 2021).

Disintegrins have served as valuable molecular templates for the development of antiplatelet and antithrombotic drugs, owing to their high specificity and affinity for integrin receptors, particularly αIIbβ3, the major and best-characterised platelet target. A notable example is eptifibatide, a cyclic heptapeptide derived from the barbourin disintegrin found in the venom of *Sistrurus miliarus barbouri*. Eptifibatide functions by mimicking the KGD (Lys-Gly-Asp) motif of barbourin, competitively inhibiting fibrinogen binding to αIIbβ3 and thereby preventing platelet aggregation (Coller, 1997; Scarborough et al., 1993). Its clinical utility in preventing thrombotic complications in patients with acute coronary syndromes underscores the therapeutic relevance of disintegrin-based drug design.

Elucidating the structure–function relationships of disintegrins is critical for the rational design of next-generation integrin-targeting therapeutics. Minor variations in amino acid sequence or loop conformation can result in dramatic shifts in integrin-binding specificity and functional activity (Calvete et al., 2005). Here, we explored the relationship between integrin-binding specificity and biological activity by expressing disintegrins in a bacterial expression system and characterising the purified, recombinant variants containing RGD, KGD, or VGD motifs. We assessed their effects on platelet aggregation and endothelial cell migration, a key aspect of angiogenesis and a useful measure of the potential of disintegrins to impair wound healing. For one RGD-containing recombinant disintegrin, we further compared its activity with a recombinant PII SVMP-derived disintegrin and the native disintegrin ocellatusin from *Echis romani*.

## 2. Materials and Methods

### 2.1. Expression of disintegrins in *E. coli*

Four disintegrins were selected for recombinant expression: EcDis_VGD (UniProt ID E9JGG6), EoDis_RGD (UniProt ID Q14FJ4), EcDis_KGD (UniProt ID E9JGE6), and EcDis_RGD (UniProt ID E9JGF7). Disintegrin constructs in the plasmid pET-24a(+) were designed and synthesised (Twist Biosciences) containing an N-terminal 6x His tag followed by DsbC (Disulfide bond C, a prokaryotic disulfide bond isomerase) (Turchetto et al., 2017) for solubility and to aid folding, and a C-terminal AviTag for biotinylation at a single site. The 6x His tag and DsbC were separated from the disintegrin by a modified TEV protease cleavage site (ENLYFQ/N) that, unlike the canonical TEV recognition sequence (ENLYFQ/G), was selected to ensure that cleavage generates the native N-terminus of the disintegrin. The plasmid with the gene of interest together with a plasmid containing the CyDisCo components (pMJS205, co-expressing PDI and Erv1p sulfhydryl oxidase) (Nguyen et al., 2011) were co-transformed into electro-competent *E. coli* BL21 (DE3) cells and then streaked out onto LB agar plates containing 34 μg/ml chloramphenicol (Sigma-Aldrich) and 50 μg/ml kanamycin (Sigma-Aldrich). A single colony was used to inoculate 10 mL LB supplemented with the aforementioned antibiotics and grown overnight at 37 **°**C, 220 rpm in a shaking incubator (New Brunswick Innova 44/44R, Eppendorf). For expression, 2.5 mL preculture was added to baffled flasks containing 250 mL of Auto Induction Media Terrific Broth Base including Trace Elements (ForMedium) supplemented with antibiotics, and incubated at 25 **°**C, 200 rpm for 24 h. Cells were pelleted at 10,000 × *g* for 10 minutes, frozen and stored at -20 **°**C until purification.

### 2.2. Purification of recombinant disintegrins

Bacterial cell pellets from 500 mL cell culture expressing the His-tagged DsbC-Disintegrin fusions were resuspended in 30 mL Lysis Buffer (50 mM Tris-HCl, 300 mM NaCl, pH 8, 0.25 mg/ml lysozyme (Sigma-Aldrich)), supplemented with 0.05% (v/v) Benzonase Nuclease (Sigma-Aldrich) and 2 mM MgSO_4_. The cells were incubated at 37 **°**C for 30 minutes and lysed by sonication for 6 minutes in cycles of 30 seconds on and 40 seconds off, with 50% amplification, while the sample was kept on ice. The cell debris were pelleted by centrifugation at 18000 × *g* for 30 minutes at 4 °C, and the cleared supernatant, containing the disintegrin DsbC His tag fusion, was filtered through a 0.22 μm syringe filter and loaded onto a 1 mL HisTrap FF column (Cytiva) pre-equilibrated with 50 mM Tris-HCl, 300 mM NaCl, pH 8, attached to an AKTA Pure system. The protein was eluted using an imidazole gradient from 0 mM to 500 mM in 50 mM Tris-HCl, 300 mM NaCl, pH 8. The elution fractions were pooled and dialysed overnight using 6-8 kDa MWCO Spectra Dialysis Membrane at 4 **°**C into 50 mM Tris, 300 mM NaCl, pH 8. To remove the DsbC tag, 0.5 mM EDTA, 0.2 mM DTT, and TEV protease (1:12 molar ratio) were added to the dialysed sample and incubated overnight at room temperature. TEV protease and uncleaved protein were removed by loading the overnight reaction onto a 1 mL HisTrap FF column (Cytiva) pre-equilibrated with 50 mM Tris, 300 mM NaCl, pH 8. The flow-through containing the disintegrin was collected, concentrated and dialysed into Low Salt buffer (20 mM Tris-HCl, 25 mM NaCl, pH 9) using an Amicon Ultra-15 Centrifugal Filter 3 kDa MWCO (Merck Millipore) and loaded onto the HiTrap Q XL 1 mL column (Cytiva). The protein was eluted using a sodium chloride gradient from 25 to 500 mM in 20 mM Tris-HCl, pH 9. Fractions containing disintegrin were pooled and concentrated to a final volume of 0.5 mL. This concentrated sample was loaded onto the Superdex 75 10/300 GL column (Cytiva) equilibrated in 50 mM Tris-HCl, 300 mM NaCl, pH 8. Samples taken from each fraction corresponding to a peak (detected at 280 nm) were analysed by reducing SDS-PAGE. Fractions containing disintegrin were pooled, concentrated and glycerol was added to 9%. The protein was flash-frozen in liquid nitrogen for long-term storage at -80 °C.

### 2.3. Isolation of ocellatusin from venom

Venom was extracted from multiple, adult, wild-caught specimens of *Echis romani* (Nigeria) (previously classified as part of *Echis ocellatus* in the broad sense (*sensu lato*)) maintained in the UK Home Office-regulated herpetarium facility at the Liverpool School of Tropical Medicine (LSTM). All chromatography was carried out using either an AKTA LC system (Cytiva) or a Vanquish HPLC system (Thermo Fisher Scientific). Buffers were freshly prepared and vacuum filtered (0.1 µm) immediately prior to use. 40 mg of freeze-dried pooled *E. romani* venom was resuspended in 2 mL ice-cold PBS (50 mM Sodium phosphate, 0.15 M NaCl pH 7.2) and centrifuged at 10,000 x *g* for 10 min. The supernatant was immediately loaded onto a 320 mL Superdex 75HR column (Cytiva) equilibrated in PBS. The column was operated at a flow rate of 1.5 mL/min and 5 mL fractions were collected after the void volume. Elution was monitored at 214 and 280 nm. SDS-PAGE analysis was carried out on all protein-containing fractions to determine which to select for the second stage of isolation. Ocellatusin eluted in a single small peak near to PLA2. The peak was detectable at 214 nm, but not at 280 nm. A Coomassie-stained band was present on the SDS-PAGE gel at around 10 kDa. To obtain a highly purified preparation, the ocellatusin-containing fractions were pooled, desalted into 50 mM Tris-HCl, pH 8.6 and applied to a 1 mL Mono-Q anion exchange column (Cytiva) equilibrated in the same buffer. Elution of ocellatusin was carried out using a 20-column volume (CV) gradient from 0 to 150 mM NaCl in 50 mM Tris-HCl, pH 8.6. The column was operated at 0.5 mL/min and 0.5 mL fractions were collected. Elution was monitored at 214 and 280 nm.

To check purity and to determine intact mass, the eluted material was added to 0.1% (v/v) formic acid (FA) and subjected to RP-HPLC-MS analysis performed on a Vanquish HPLC system coupled to an Orbitrap Exploris 240 mass spectrometer (Thermo Fisher Scientific).

RP-HPLC separation was carried out on a Hypersil Gold C18 column (175 Å pore size, 100 × 2.1 mm, 1.9 μm particle size) with the column oven set to 50°C. The flow rate was 0.3 mL/min and the proteins were eluted with a gradient of 0.1% formic acid (FA) in water (buffer A) and 0.1% FA in acetonitrile (ACN) (buffer B): 2–40% B for 0–10 min, 40–80% B for 10–12 min, 80% B for 12–15 min, before returning to 2% B at 15–15.1 min and re-equilibrating at 2% B for 15.1–20 min. ESI settings were as follows: sheath gas, 5 arbitrary units (Arb); auxiliary gas, 2 Arb; spray voltage, 3.5 kV; ion transfer tube temperature, 275 °C; mass tolerance, 15 ppm. The Orbitrap full scan acquisition parameters were: resolution, 120,000 (at m/z 200); scan range, m/z 200–2,000; radio frequency lens (%) set to 70, polarity positive with the automatic gain control (AGC) target set to standard.

### 2.4. PII SVMP production

EoSVMP_PII_zym_ expression and purification were performed as described previously (Hall et al., 2026). Briefly, the protein was expressed in baculovirus infected Hi5 insect cells at 19°C, in ESF 921 Insect Cell Culture Media (Expression Systems). Cultures were harvested 5 days after infection by centrifugation (1000 x *g*, 10 minutes), and the supernatant was passed through HiTrap IMAC FF (Cytiva), followed by purification by ion exchange chromatography using a HiTrap Q XL column (Cytiva). Finally, the SVMP was purified by size exclusion chromatography using a Superdex 200 increase 10/300 GL column (Cytiva). Eluted PII SVMP was concentrated, and glycerol was added to 5% before the protein was flash-frozen in liquid nitrogen and stored at -80 °C till use. To generate the mature, autoactivated SVMP PII (EoSVMP_PII), EoSVMP_PII_zym_ was incubated with 750 μM ZnCl_2_ overnight at 37 °C.

To determine the intact mass of the product formed by autoactivation, EoSVMP_PII was subjected to mass spectrometry analysis, as follows: 5 μL of concentrated sample solution was transferred to a new tube and diluted to a final volume of 20 µL prior to desalting using C4 ZipTips (Millipore). Following desalting, the samples were dried using a SpeedVac SPD121P vacuum concentrator (Thermo Fisher Scientific). The dried samples were then resuspended in 10 µL of a solution containing acetonitrile, ultrapure water, and formic acid (50/49.9/0.1, v/v/v). In addition, a 2 mg/mL bovine serum albumin (BSA) standard solution (Sigma-Aldrich) was used as a calibrant. The BSA standard was desalted following the same procedure as the samples and subsequently resuspended in 2 µL of the acetonitrile/ultrapure water/formic acid solution described above. Samples and calibrant were deposited onto the MALDI target using the dried-droplet method. Briefly, 1 µL of sample or calibrant was spotted onto the target plate, followed by the addition of 1 µL of sinapinic acid (SA) matrix solution. The spots were allowed to dry at room temperature before analysis. Mass spectra were acquired using a rapifleX® MALDI-TOF/TOF mass spectrometer (Bruker Daltonics, Bremen, Germany) in positive-ion, linear mode over an m/z range corresponding to 2–80 kDa.

### 2.5. Plate aggregometry

This research involving human blood samples has been reviewed and approved by the research ethics committee of the United Bristol Healthcare NHS Trust (REC reference 20/SC/0222; ’Studies of human platelet functional and signalling events’). Human blood was obtained from healthy, drug-free volunteers who provided full informed consent in accordance with the Declaration of Helsinki. Blood was drawn via venepuncture from healthy volunteers into BD vacutainers containing 3.2% sodium citrate. Inclusion criteria were adult volunteers aged 18 or over at the time of enrolment. Donors were excluded if they had a known clotting or bleeding disorder, regularly consumed medication that may impair platelet function (i.e., aspirin, ibuprofen), were pregnant or had a known blood-borne disease.

Blood was centrifuged at 180 x *g* for 17 min at room temperature to recover platelet-rich plasma (PRP) and centrifuged at 550 x *g* for 10 min to recover platelet-poor plasma (PPP). Agonists were prepared in HEPES-Tyrode’s buffer (10 mM HEPES pH 7.3, 145 mM NaCl, 1 mM MgSO_4_, 3 mM KCl). Platelet aggregation experiments were performed using an end-point assay in 96-well half-area plates. Disintegrins (5 µL) were incubated with 45 μL PRP for 5 min at 37 °C prior to platelet aggregation induction by 2-MeS-ADP (0.25 nM to 5 μM) (Tocris Bioscience), CRP-XL (0.25 nM to 5 μg/mL) (Triple Helical Peptides), or TRAP-6 (1 nM to 20 μM) (Bachem). The plate was shaken at 1200 rpm, 37 °C, and after 5 min, absorbance was read at 595 nm using the LT4500 plate reader (Labtech). To determine % aggregation, the sample absorbance was normalised, accounting for unstimulated PRP as 0% aggregation and PPP sample absorbance as 100% aggregation. For the disintegrins concentration-response curves at fixed agonist concentration, the absorbance value of the control (without disintegrins) was normalised to 100% of aggregation. All mean values and IC_50_s were calculated based on three biological replicates performed in duplicate. Data were plotted, and E_max_ was calculated using a four-parameter variable slope model (log[agonist] vs. response), with the bottom constraint set to zero in GraphPad Prism version 11 (GraphPad Software).

### 2.6. Scratch wound healing assay

HUVEC cells (PromoCell) were seeded into a 96-well ImageLock tissue culture plate (Incucyte) at 20,000 cells per well in 100 µL complete medium (Endothelial Cell Growth Medium 2 with supplements, PromoCell) and incubated for 24 h at 37 °C, 5% CO₂ to achieve a confluent monolayer. Cells were then treated with 10 μg/mL mitomycin C (Sigma-Aldrich) in serum-free medium (Endothelial Cell Growth Basal Medium 2, PromoCell) for 2 h. Straight scratches were made with a 96-pin wounding tool through the centre of each well. Each well was gently washed with 100 μL PBS to remove detached cells. 100 µL complete medium containing disintegrins at the desired concentrations (1000, 500 and 250 nM) was added to the experimental wells. 100 μL complete medium with buffer (50 mM Tris-HCl, 300 mM NaCl, pH 8) was used as an untreated scratch control. Cells were incubated under standard conditions (37 °C, 5% CO₂) in the Incucyte Live-Cell Analysis System (Sartorius), and phase-contrast images were acquired at time 0, after 2 h, and then every hour until 23 h, using a 10x objective to monitor wound closure. Image acquisition and analysis were performed using Incucyte software version 2022A, applying the integrated wound analysis module. Cell migration was quantified using the Relative Wound Density (RWD), which measures the density of cells within the wound area relative to the density outside the wound region, normalised to the initial time point. RWD values were automatically calculated by the software for each time point. Data were expressed as percentage wound closure relative to the initial wound (time 0). For each condition, two technical replicates per experiment and three independent biological repeats were performed. Data were plotted as RWD (%) over time and as a bar graph with the corresponding cell migration calculated as the % of RWD normalised to the control after 23 h. Statistical analyses were conducted using Two-way ANOVA with post hoc Dunnett’s multiple comparisons test in GraphPad Prism version 11.

### 2.7. MTT cell viability assay

HUVEC cells were seeded as above and incubated for 24 h at 37 °C, 5% CO₂, to achieve a confluent monolayer. The cell line was freshly procured and checked for contamination before the start of the experiment. The next day, cells were treated with 10 μg/mL mitomycin-C for 2 h, before adding 100 µl/well (duplicate wells) of the recombinant disintegrin treatments: 250, 500 and 1000 nM prepared in Endothelial Cell Growth Medium 2. After 24 h, 10 μl of MTT solution from the MTT Cell Viability Assay Kit (Biotium) was added to the 100 μl of medium in each well and mixed by tapping gently on the side of the tray. The cells were incubated at 37 °C for 4 h. 200 μl DMSO was added directly to the medium in each well and pipetted up and down several times to dissolve the formazan salt. The absorbance was measured on BioTek Synergy Neo2 spectrophotometer at 570 nm. The background absorbance was also measured at 630 nm (subtracted background absorbance from signal absorbance to obtain normalised absorbance values). The % of cell viability for each treatment well was calculated as follows, with buffer-treated cells as a control (100% viability):

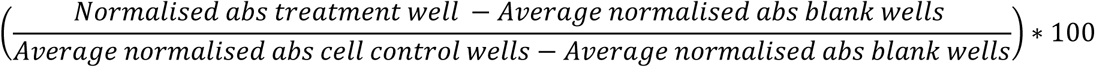

## 3. Results

### 3.1. Production of *Echis* disintegrins in *E. coli*

We designed DNA constructs for the expression of disintegrins from *E. coloratus and E. ocellatus* containing distinct integrin-binding motifs: EcDis_VGD (UniProt ID E9JGG6; VGD motif), EcDis_KGD (UniProt ID E9JGE6; KGD motif), EcDis_RGD (UniProt ID E9JGF7; RGD motif) and EoDis_RGD (UniProt ID Q14FJ4; RGD motif) (Figure 1A). The disintegrins share moderate to high sequence identity: EcDis_RGD and EoDis_RGD are the most closely related pair (87.7%), whereas EcDis_RGD and EcDis_KGD show the lowest identity (56.0%). EcDis_VGD shares 69.7% identity with EcDis_KGD, 72.8% with EcDis_RGD, and 74.1% with EoDis_RGD, while EoDis_RGD shares 59.1% identity with EcDis_KGD. EoDis_RGD corresponds to 100% sequence identity to ocellatusin – a well-known RGD-disintegrin from *E. ocellatus* (Figure 1D). Apart from EcDis_VGD, all other disintegrins correspond to the disintegrin domain sequence of the corresponding PII SVMPs, which comprise a metalloproteinase and a disintegrin domain. According to the ADDovenom toxin database (Redureau et al., 2025), these four disintegrins and their corresponding SVMPs showed variable relative abundances in the venoms, with EcDis_VGD (2.82%) and EcDis_KGD (2.97%) being most abundant, followed by EoDis_RGD (2.27%) and EcDis_RGD (0.24%).

**Figure 1.**
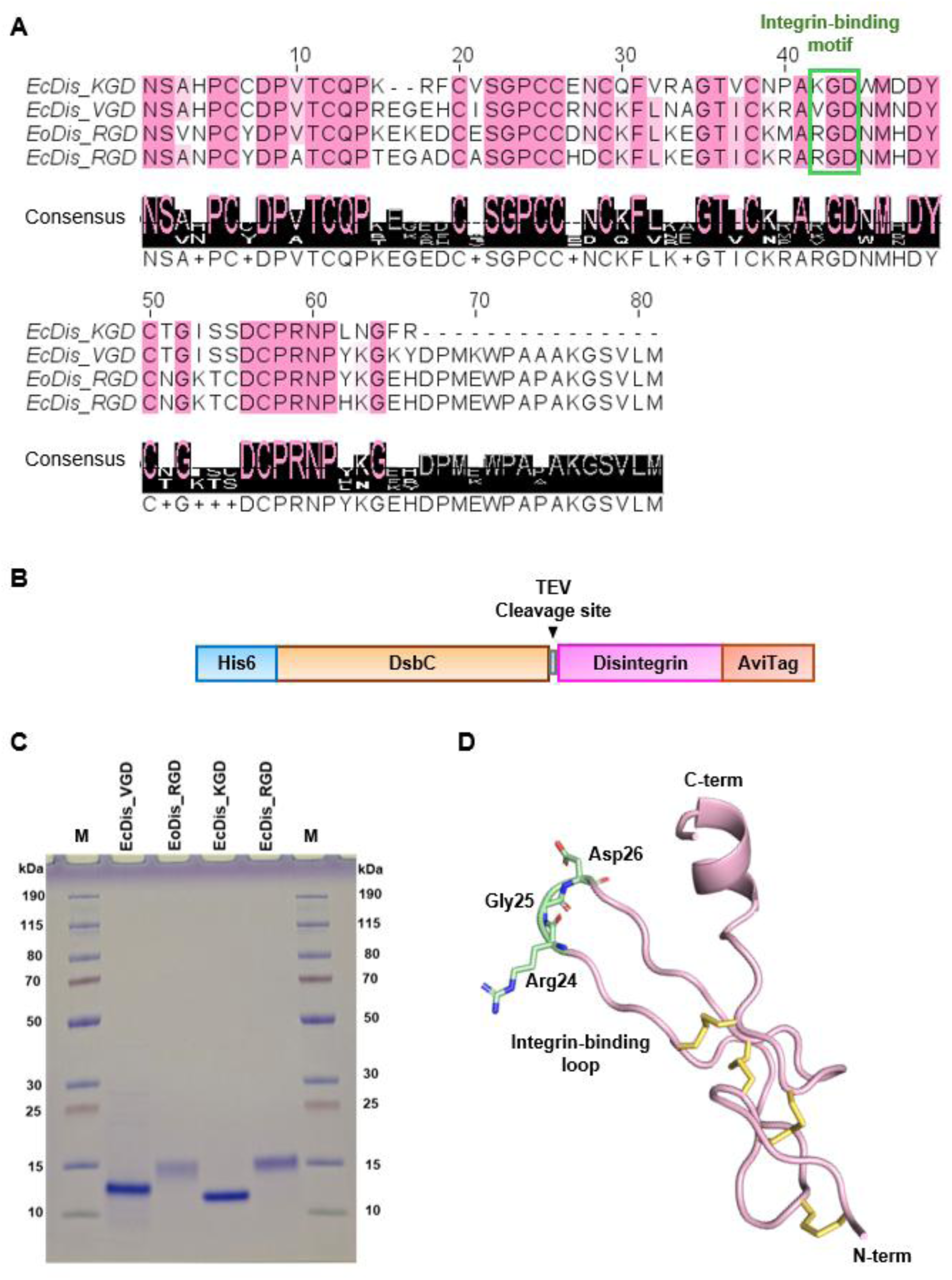
Sequence analysis, construct design, purification of recombinant disintegrins and disintegrin structural model. **A)** Sequence alignment of EcDis_KGD (UniProt ID E9JGE6), EcDis_VGD (UniProt ID E9JGG6), EoDis_RGD (UniProt ID Q14FJ4), and EcDis_RGD (UniProt ID E9JGF7). The integrin-binding motifs are highlighted in a green box. Multiple sequence alignment was generated by Clustal Omega and visualised using Jalview. **B)** Schematic representation of the recombinant disintegrin construct design containing an N-terminal 6x His tag, DsbC (Disulfide bond C - a prokaryotic disulfide bond isomerase), a modified TEV protease cleavage site (ENLYFQ/N), the disintegrin, and a C-terminal AviTag. **C)** Coomassie-stained reducing SDS-PAGE of purified Ec_VGD, Eo_RGD, Ec_KGD, and Ec_RGD, following size-exclusion chromatography. M: PageRuler™ Plus Prestained Protein Ladder. **D)** AlphaFold3 structural model of Ocellatusin (UniProt ID Q3BER1). Disulfide bonds are coloured in yellow. The RGD motif is highlighted in green. C- and N-termini are indicated.

Our expression constructs consist of an N-terminal 6x His tag followed by DsbC and the gene encoding the disintegrin. DsbC is a periplasmic disulfide-bond isomerase and molecular chaperone that promotes the correct formation and rearrangement of disulfide bonds during protein folding (Turchetto et al., 2017). The disintegrin is followed by a C-terminal AviTag for biotinylation (Figure 1B). His-tag and DsbC can be separated from the disintegrin by TEV protease cleavage. A modified TEV cleavage site was used to provide the disintegrins with their native N-terminus (asparagine).

The disintegrins were successfully expressed in *E. coli* BL21(DE3) containing the plasmid pMJS205 used for co-expressing protein disulfide isomerase (PDI) and Erv1p sulfhydryl oxidase, to further promote the formation and isomerisation of disulfide bonds (Sohail et al., 2020). The disintegrin purification protocol included immobilised metal affinity (IMAC), anion exchange (AIEX) and size exclusion chromatography (SEC) (Figure S1). To remove the DsbC tag, TEV proteinase cleavage was performed before the AIEX step. The disintegrins were obtained as monomeric proteins with high purity and satisfactory yields of 0.3, 0.6, 0.8, and 1.2 mg per litre for EcDis_VGD, EcDis_KGD, EcDis_RGD and EoDis_RGD, respectively (Figure 1C, S1 and S2).

### 3.2. Evaluation of recombinant disintegrins on human-platelet activity

To assess the effects of disintegrins on platelet aggregation, we generated concentration– response curves using three agonists that promote platelet aggregation by targeting different receptors: 2-MeS-ADP (2-methylthioadenosine diphosphate), a potent P2Y1 and P2Y12 receptor agonist (Gachet, 2006); CRP-XL (cross-linked collagen-related peptide), an activator of the glycoprotein VI (GPVI) receptor (Smethurst et al., 2007); and TRAP-6 (thrombin receptor-activating peptide 6), a selective protease-activated receptor 1 (PAR1) agonist (Zou et al., 2024). Assays were performed in the presence and absence of disintegrins using endpoint plate aggregometry.

When platelet aggregation responses to 2-MeS-ADP were assessed in the presence of 50 nM EoDis_RGD and EcDis_RGD, the concentration-response curves were shifted to the right relative to the control, suggesting an interference during the activation phase, while the maximal response (E_max_) remained unchanged at 98.2% and 97.6% for EoDis_RGD and EcDis_RGD, respectively (Figure 2A, Table 1). At higher RGD motif-containing disintegrin concentrations (250 nM), aggregation was markedly attenuated across the 2-MeS-ADP concentration range used, with low responses observed even at the highest agonist concentrations (Table 1). Platelet aggregation was completely abolished at 500 nM disintegrin concentrations, with responses remaining at baseline levels irrespective of agonist concentration (Figure 2A). In contrast, EcDis_KGD at 50 nM showed no effect, while at 250 nM, the concentration-response curve shifted to the right relative to the control. At 500 nM disintegrin concentration, aggregation was reduced, and the maximal response E_max_ reached only 60% (Figure 2A, Table 1).

**Figure 2.**
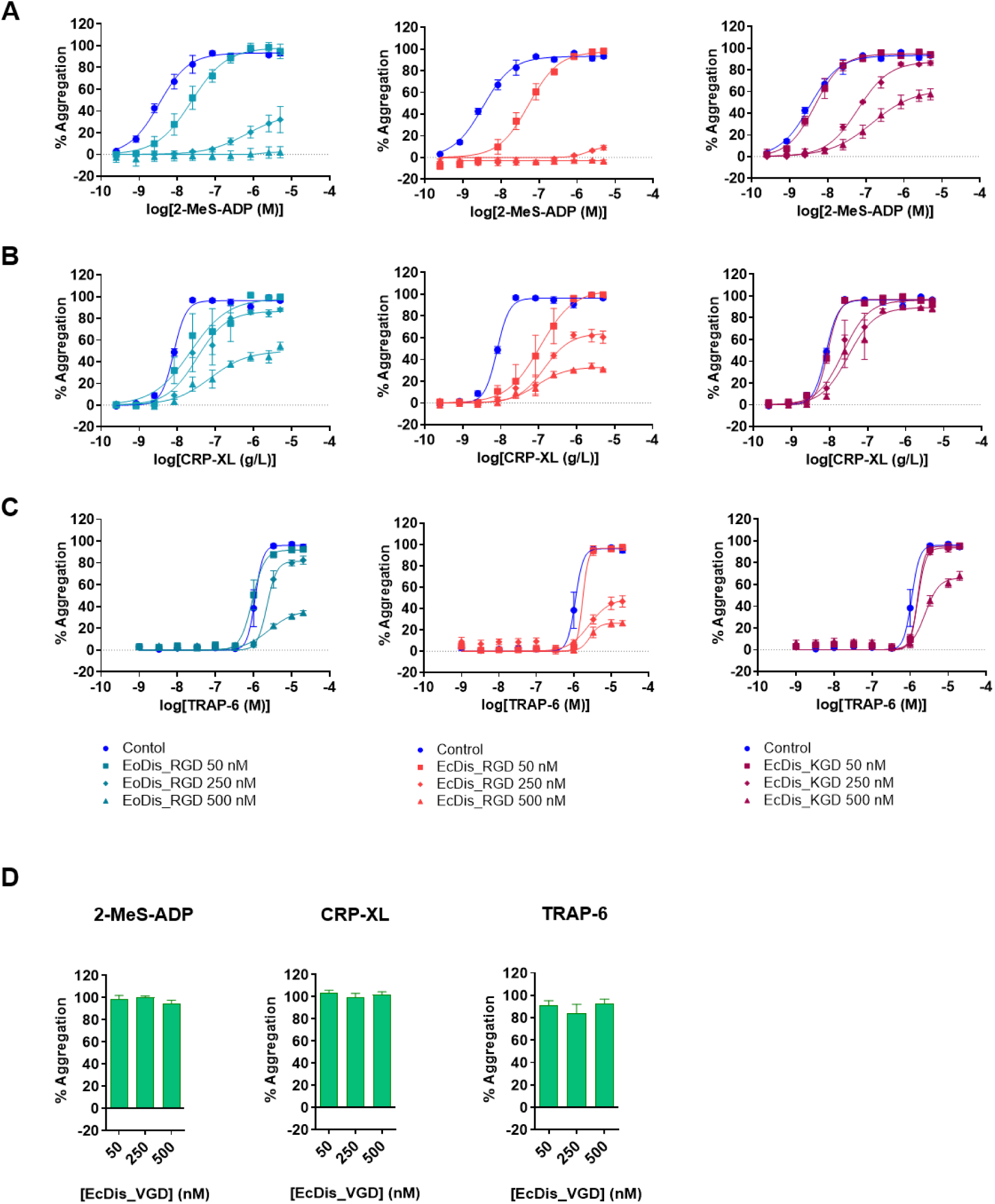
Effect of disintegrins on concentration-dependent platelet aggregation responses to **A)** 2-MeS-ADP, **B)** CRP-XL, and **C)** TRAP-6. Control curves (without disintegrins) are shown in blue circles. Disintegrin-treated samples were tested at 50 nM (squares), 250 nM (diamonds), and 500 nM (triangles). EcDis_KGD (magenta curves), EoDis_RGD (dark teal curves) and EcDis_RGD (red curves). PRP was pre-incubated with disintegrins for 5 min before stimulation with increasing concentrations of agonist. After 5 min of agonist stimulation, absorbance was measured at 595 nm. Data are presented as the mean of % aggregation ± SEM versus agonist concentration. Curves were fitted using a four-parameter variable slope model (log[agonist] vs. response), with the bottom constraint set to zero, in GraphPad Prism 11. **D)** Bar graph showing the percentage of aggregation with 2.5 μM 2-MeS-ADP, 2.5 μg/mL CRP-XL and 3.3 μM TRAP-6, in the presence of EcDis_VGD. The assays were carried out in experimental duplicate and biological triplicate. Data are presented as the mean of % aggregation ± SEM.

**Table 1.**
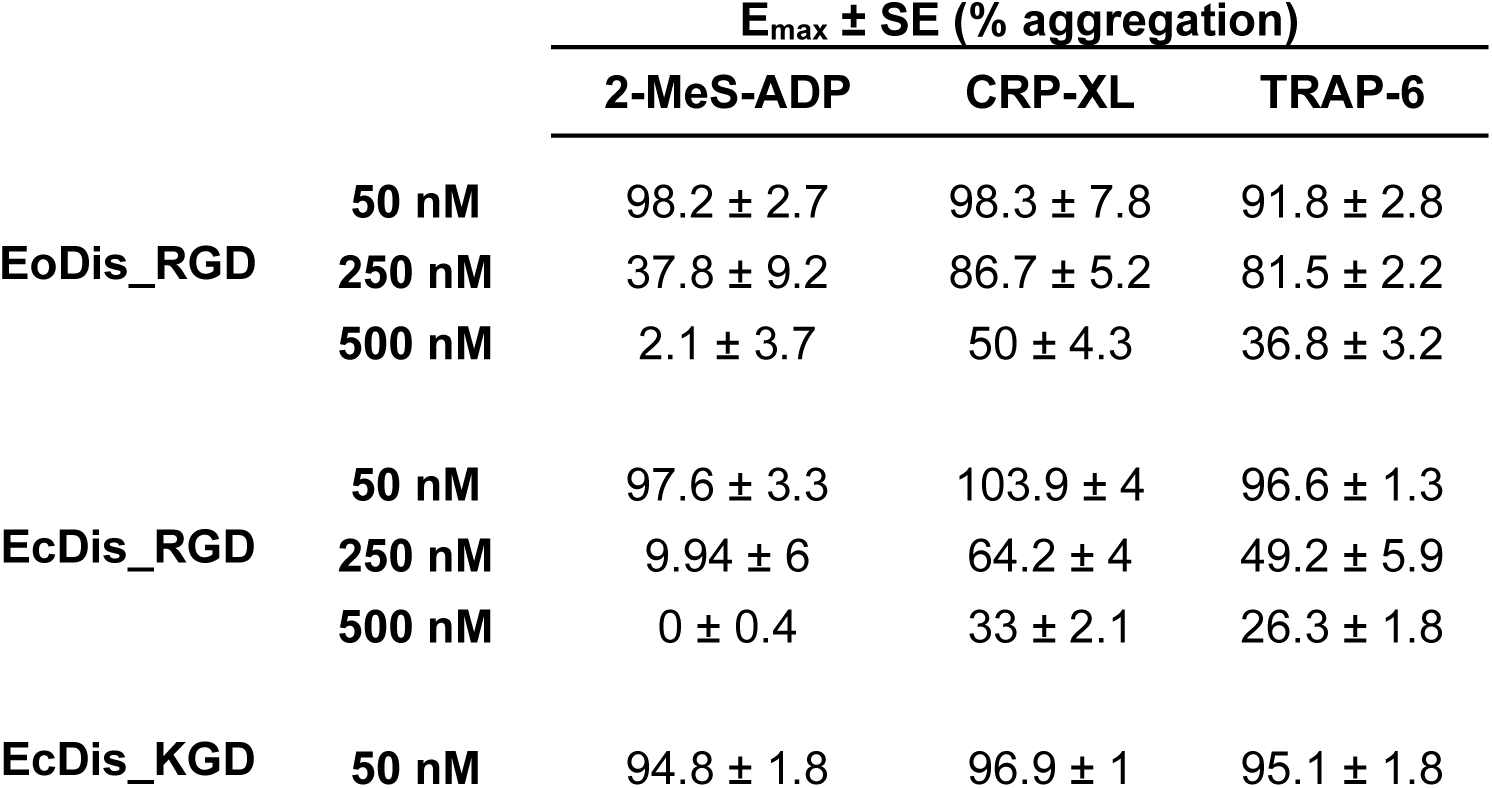
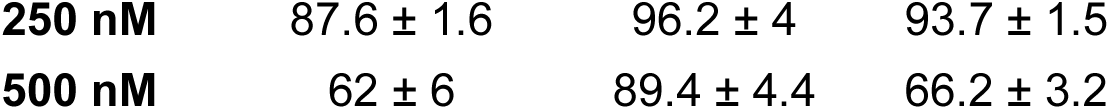
Maximum response (E_max_) values (% aggregation ± SE) for 2-MeS-ADP-, CRP-XL-, and TRAP-6-induced platelet aggregation in the presence of indicated concentrations of recombinant disintegrins.

Similarly, when CRP-XL was assessed in the presence of 50 nM EoDis_RGD or EcDis_RGD, a rightward shift of the concentration-response curves was observed, while the E_max_ remained the same (Figure 2B). At 250 nM, EoDis_RGD shifted the curve and reduced E_max_ slightly (86.7%), while EcDis_RGD presented a much lower E_max_ (64.2%). At the highest disintegrin concentration (500 nM), EoDis_RGD diminished platelet aggregation to 50%, while EcDis_RGD reduced it to 33% (Table 1). In contrast, EcDis_KGD showed no detectable effect at 50 nM. However, at concentrations of 250 and 500 nM, a rightward shift in the concentration-response curve was observed, with the highest concentration reducing E_max_ to 89.4% (Figure 2B, Table 1).

Using TRAP-6 as the agonist, the disintegrins exhibited minimal effects on platelet aggregation at 50 nM (Figure 2C). At 250 nM, EoDis_RGD and EcDis_RGD reduced the E_max_ to 81.5% and 49.2%, respectively. Increasing the concentration to 500 nM further increased inhibition, with E_max_ values decreasing to 36.8% for EoDis_RGD and 26.3% for EcDis_RGD (Table 1). In contrast, EcDis_KGD affected platelet aggregation only at 500 nM, reducing the E_max_ to 66.2% (Table 1).

Unlike the other recombinant disintegrins, EcDis_VGD did not affect platelet aggregation at any of the tested concentrations (50, 250 or 500 nM), regardless of the agonist used (2-MeS-ADP, CRP-XL, or TRAP-6) (Figure 2D).

In summary, RGD-containing disintegrins, and to a lesser extent the KGD-containing disintegrin, produced concentration-dependent inhibition of platelet aggregation, characterised by progressive attenuation of the agonist responses and a marked reduction in maximal aggregation at higher concentrations.

For all agonists tested, EcDis_RGD exhibited the strongest inhibitory profile. We determined half-maximal inhibitory concentrations (IC_50_s) of 144 nM (95% CI: 127 – 163), 292 nM (95% confidence interval (CI): 268 – 317), and 235 nM (95% CI: 208 – 265) for 2-MeS-ADP, CRP-XL and TRAP-6, respectively. EoDis_RGD had IC_50_ values of 186 nM (95% CI: 172 – 202), 394 nM (95% CI: 359 – 432), and 268 nM (95% CI: 195 – 367) for 2-MeS-ADP, CRP-XL and TRAP-6, respectively. EcDis_KGD was the least potent inhibitor, with IC_50_ values of 411 nM (95% CI: 344 – 491), 650 nM (95% CI: 567 – 745), and 597 nM (95% CI: 469 – 760) for 2-MeS-ADP, CRP-XL and TRAP-6, respectively (Figure 3, Table 2). Notably, although EoDis_RGD, EcDis_RGD, and EcDis_KGD produced concentration-dependent inhibition, all curves reached a plateau at residual aggregation levels of approximately 20% when the agonists CRP-XL and TRAP-6 were used, indicating that a fraction of the platelet response remained unaffected even at the highest disintegrin concentrations analysed (Figure 3).

**Figure 3.**
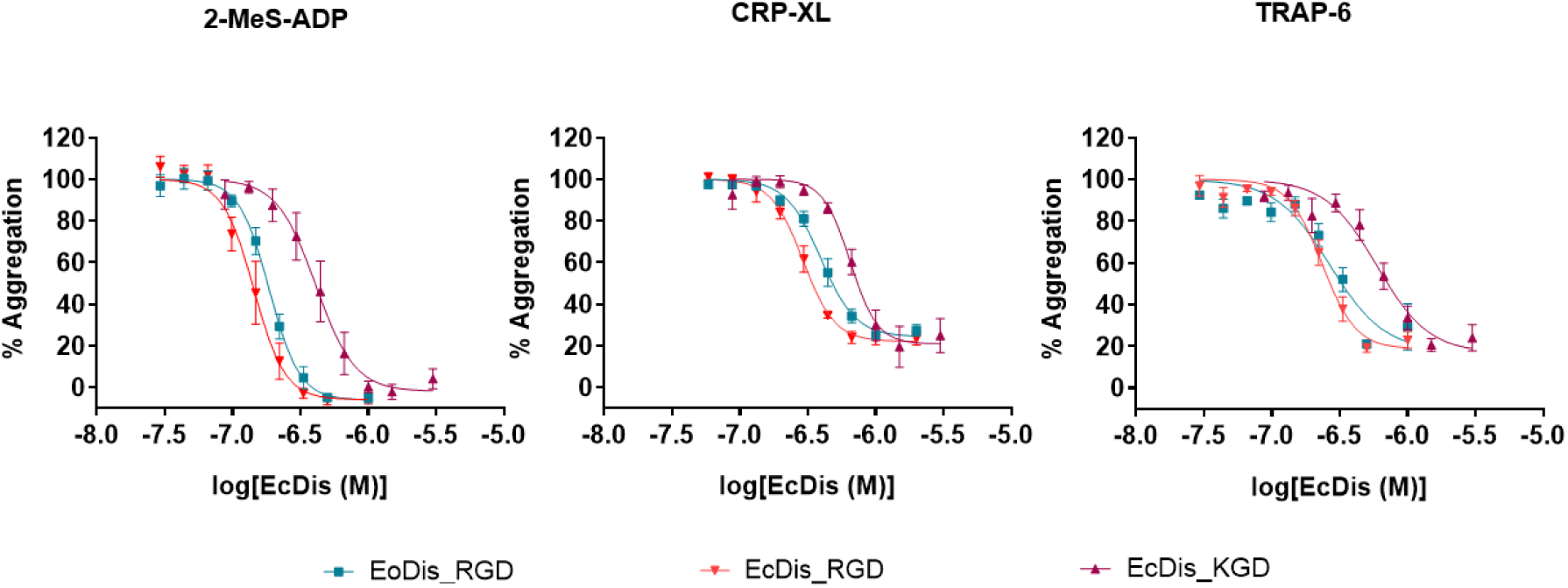
Concentration-response curves for IC50 determination of EcDis_KGD, EoDis_RGD and EcDis_RGD against 2-MeS-ADP-, CRP-XL- and TRAP-6-induced platelet aggregation. Human platelet-rich plasma was pre-incubated for 5 min with varying concentrations of EcDis_KGD, EoDis_RGD and EcDis_RGD, followed by stimulation with 2.5 μM 2-MeS-ADP, 2.5 μg/mL CRP-XL or 3.3 μM TRAP-6. EcDis_KGD (magenta), EoDis_RGD (dark teal) and EcDis_RGD (red). The assays were carried out in experimental duplicate and biological triplicate. Data are presented as the mean of % aggregation ± SEM versus disintegrin concentration. Concentration-response curves were fitted using a four-parameter logistic model (log[inhibitor] vs. response – variable slope), with the top constraint set to one hundred, in GraphPad Prism 11.

**Table 2.**
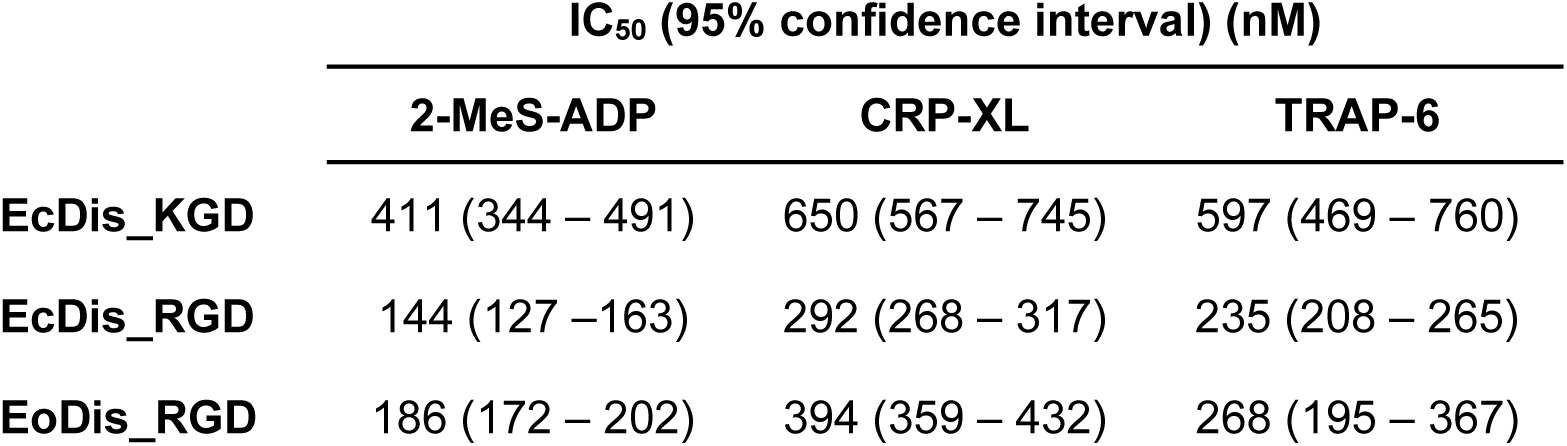
IC_50_ values for recombinant disintegrin-mediated inhibition of 2-MeS-ADP-, CRP-XL- , and TRAP-6-induced platelet aggregation, expressed in nM.

|  | IC<sub>50</sub> (95% confidence interval) (nM) |  |  |
| --- | --- | --- | --- |
|  | 2-MeS-ADP | CRP-XL | TRAP-6 |
| **EcDis_KGD** | 411 (344 – 491) | 650 (567 – 745) | 597 (469 – 760) |
| **EcDis_RGD** | 144 (127 – 163) | 292 (268 – 317) | 235 (208 – 265) |
| **EoDis_RGD** | 186 (172 – 202) | 394 (359 – 432) | 268 (195 – 367) |

### 3.3. Effect of recombinant disintegrins on human endothelial cell migration

A scratch wound healing assay was performed using human umbilical vein endothelial cells (HUVEC) to evaluate the ability of each disintegrin to inhibit cell migration. In a first approach, the viability of HUVEC after treatment with the disintegrins was analysed using MTT assays. Importantly, HUVEC viability was maintained in cells incubated with increasing concentrations of disintegrins (250 to 1000 nM) after 24 h (Figure S3).

For the scratch assay, a uniform wound was generated on a confluent HUVEC monolayer pre-treated with mitomycin C to inhibit cell proliferation. The rate of wound closure was monitored for 23 h in the presence or absence of the disintegrins (Figure 4A). During the first 5 h, wound closure progressed similarly across all conditions, but differences became more apparent at later time points (Figure S4). The strongest inhibitory effect was observed in the cells treated with EcDis_RGD; at all concentrations tested, cell migration was markedly impaired throughout the experiment, resulting in a final relative wound density of approximately 50% (250 to 1000 nM; *p <* 0.0001). In contrast, cells treated with 250 and 500 nM EoDis_RGD exhibited wound closure profiles comparable to the control group (no disintegrin), reaching approximately 90-100% relative wound density after 23 h. However, treatment with 1 µM reduced wound closure, reaching approximately 60% relative wound density at the end of the assay (*p <* 0.0001). EcDis_VGD and EcDis_KGD had no significant effect on cell migration, with migration levels remaining comparable to the control group at all concentrations tested (Figure 4B).

**Figure 4.**
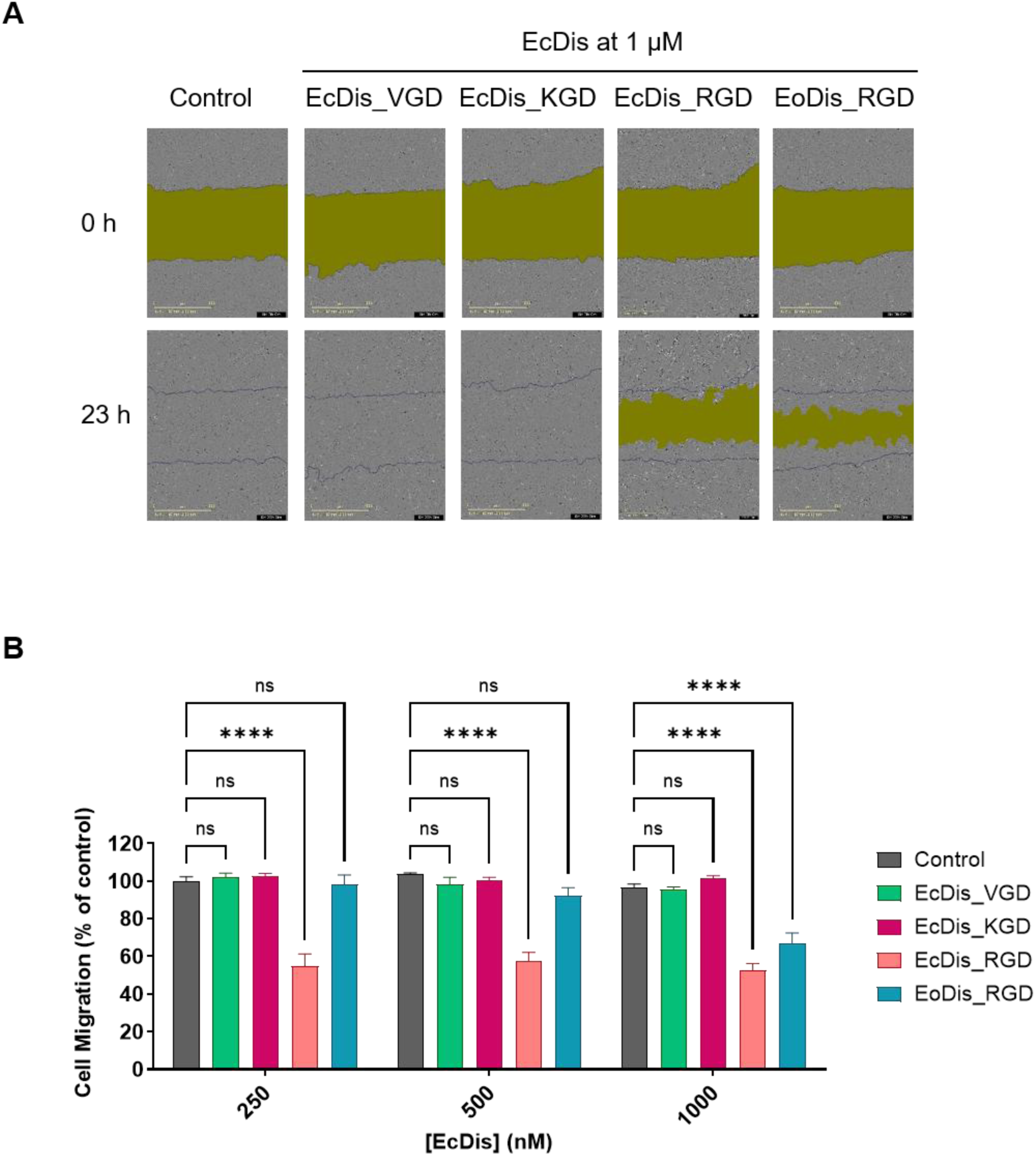
Effect of disintegrins on cell migration. **A)** Representative images of the scratch wound healing test. HUVEC were grown to confluence, after which a scratch was made across the centre of each well. Images were acquired after 23 h of disintegrin treatment using the Incucyte system. The assays were carried out in duplicate in three independent tests. **B)** Bar graph shows the percentage of migration relative to the control (buffer). Control is shown in dark grey, EcDis_VGD in green, EcDis_KGD in magenta, EcDis_RGD in red and EoDis_RGD in dark teal. Statistical analysis was performed using Two-way ANOVA and Dunnett’s Multiple Comparison post-test, in GraphPad Prism 11. \*\*\*\**p <* 0.0001.

In summary, our results indicate that RGD-containing disintegrin variants effectively impair HUVEC migration, with EcDis_RGD presenting a more potent inhibitory effect than EoDis_RGD, while the KGD- and VGD-containing disintegrins did not significantly inhibit wound closure under the tested conditions.

### 3.4. Comparison of the inhibitory potency of native ocellatusin with *E. coli* produced disintegrin and an SVMP PII-derived disintegrin

Ocellatusin is the major secreted disintegrin in *E. ocellatus* venom, representing around 3.9% of the venom composition (Juárez et al., 2006; Wagstaff et al., 2009). It can be generated in the venom either by proteolytic processing of a P-II snake venom metalloproteinase (EoSVMP_PII), which releases the disintegrin domain from the metalloproteinase domain, or by direct translation of a disintegrin-coding transcript lacking the metalloprotease domain and presenting the same sequence as the disintegrin domain in EoSVMP_PII (Juárez et al., 2006). The latter was expressed in *E. coli* as the recombinant disintegrin EoDis_RGD in this study. Our recombinant toxin EoDis_RGD comprises a disintegrin core domain that is identical to that of ocellatusin (Figure 5A).

**Figure 5.**
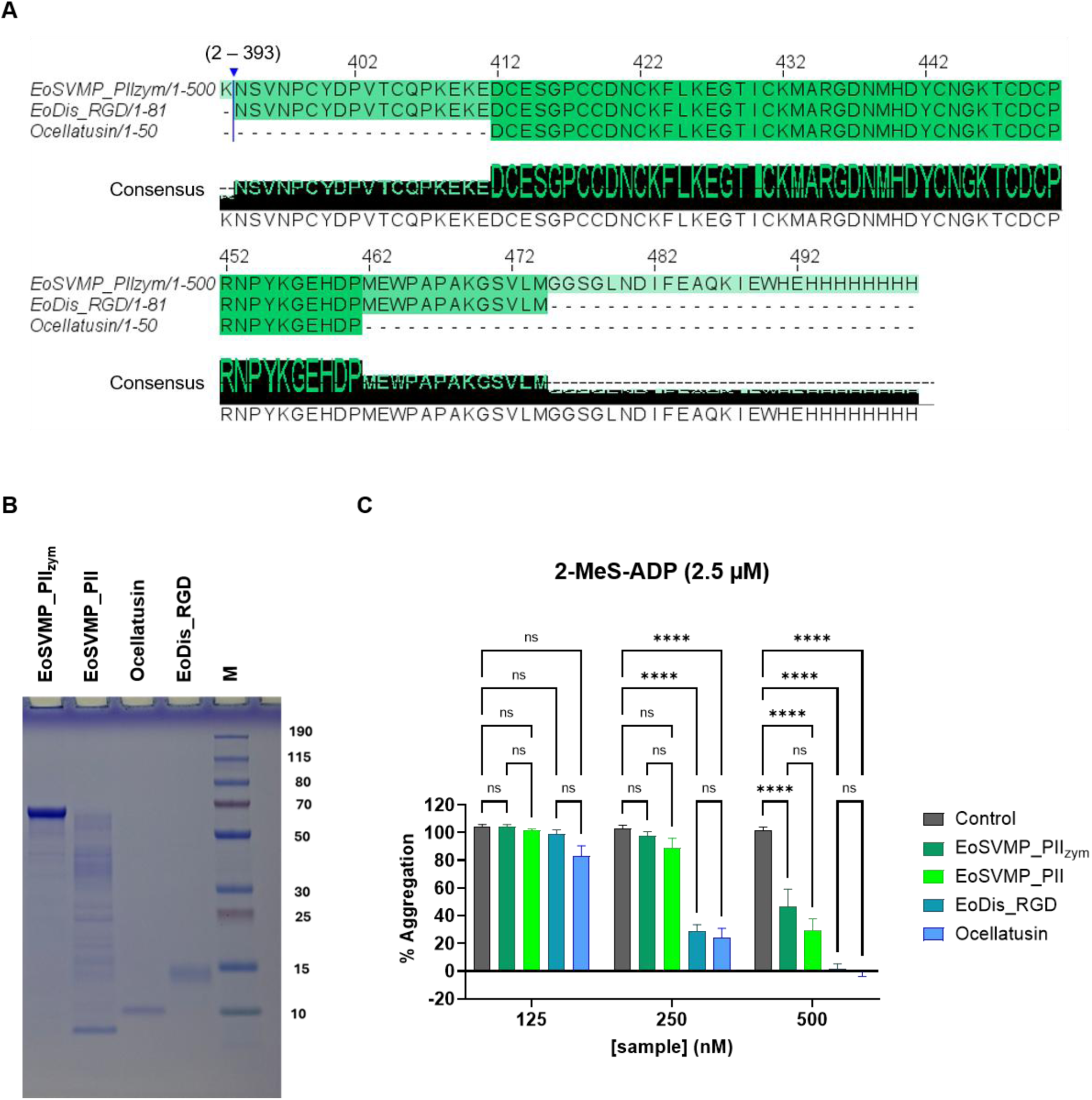
Comparison of native ocellatusin with EoDis_RGD and EoSVMP_PII. **A)** Sequence alignment of EoSVMP_PII_zym_ (UniProt ID Q14FJ4), EoDis_RGD and Ocellatusin (UniProt ID Q3BER1). EoSVMP_PII_zym_ residues from 2 to 393 are hidden for clarity purposes. Multiple sequence alignment was generated by Clustal Omega and visualised using Jalview. **B)** Coomassie-stained reducing SDS-PAGE analysis showing the EoSVMP_PII_zym_, EoSVMP_PII, venom-purified ocellatusin, and EoDis_RGD. M: PageRuler™ Plus Prestained Protein Ladder. **C)** Bar graph showing the percentage of aggregation in the presence of 2.5 μM 2-MeS-ADP. EoSVMP_PII__zym_ is shown in dark green, EoSVMP_PII is shown in light green, EoDis_RGD is shown in dark teal. The assays were carried out in experimental duplicate and biological triplicate. Data are presented as the mean of % aggregation ± SEM versus agonist concentration. Statistical analysis was obtained using Two-way ANOVA and Tukey’s Multiple Comparison post-test, in GraphPad Prism 11. ****p < 0.0001.

To understand the impact of toxin maturation on the ability to inhibit platelet aggregation, we compared the corresponding SVMP PII zymogen (inactive enzyme) and the mature, auto-activated SVMP PII (EoSVMP_PII_zym_ and EoSVMP_PII), the native ocellatusin, and the *E. coli*-produced recombinant disintegrin EoDis_RGD. EoSVMP_PII_zym_ was produced recombinantly using the MultiBac baculovirus/insect cell system and purified through IMAC, AIEX and SEC, as previously reported (Hall et al., 2026). Ocellatusin was purified from venom using SEC and AIEX (Figure S5A), and its identity was confirmed by RP-HPLC-MS analysis, which yielded three mass values: 5589.3, 5605.3, and 5621.2 Da, representing the native mass, and masses with one and two oxidised methionines, respectively (Figure S5B-E). To auto-activate EoSVMP_PII_zym_, the protein was incubated overnight in the presence of Zn^2+^ ions. In the activated SVMP (EoSVMP_PII), the disintegrin domain is released from the metalloproteinase domain after the auto-inhibitory prodomain has been cleaved off (Hall et al., 2026) (Figure 5B and S6).

All toxins tested inhibited platelet aggregation in a concentration-dependent manner. Native ocellatusin exhibited a strong inhibitory profile, similar to EoDis_RGD, with a significant reduction of platelet aggregation at 250 and 500 nM (∼25% and 0% aggregation, respectively) (*p <* 0.0001). In contrast, EoSVMP_PII_zym_ and EoSVMP_PII showed a weaker inhibitory effect, with comparable levels of inhibition at 500 nM (∼40% aggregation) (*p* < 0.0001) (Figure 5C). To better understand the differences between PII-derived disintegrin and ocellatusin, we determined the protein mass of EoSVMP_PII by matrix-assisted laser desorption/ionisation mass spectrometry (MALDI-MS) (Figure S6A). We obtained a major peak at 9005.7 Da, compared to ocellatusin, which has a molecular mass of 5621.2 Da (Smith et al., 2002). The most probable sequence with this mass has the same N-terminus as ocellatusin and comprises the C-terminal part of the PII-disintegrin domain with the AviTag but lacking the His-tag (Figure S6B). The resulting protein would have a similar molecular weight as EoDis_RGD, which is fully active and runs at an apparent MW of 14 kDa in the reducing Coomassie-stained SDS gel (Figure 5B). We note, however, that there is only a relatively faint band on the reducing Coomassie-stained SDS gel at this molecular weight (∼14 kDa) for the EoSVMP_PII sample (Figure 5A), which could explain the reduced activity in platelet aggregation assays.

Taken together, the *E. coli* derived disintegrin has comparable anti-platelet aggregation activity to the native disintegrin ocellatusin, while the disintegrin sample obtained from the PII SVMP zymogen may require more efficient folding and/or proteolytic processing to reach full activity in our experiments.

## 4. Discussion

Transcriptomic analysis of the *Echis coloratus* venom gland previously identified a cluster of short coding disintegrins that represents approximately 5% of toxin-encoding transcripts (Casewell et al., 2009). Consistent with these findings, subsequent proteomic analysis detected disintegrins in the venom, where they accounted for 0.24 to 7.7% of the total venom composition (Avella et al., 2025; Casewell et al., 2014). Disintegrins are of particular interest because they target integrins, a family of adhesion receptors that regulate platelet aggregation, cell migration, angiogenesis, inflammation, and tumour progression (Cesar et al., 2019). Small differences in disintegrin sequence, especially within their integrin-binding motifs, can markedly alter receptor selectivity and consequently their biological activities (Almeida et al., 2023). Venoms frequently contain multiple disintegrin isoforms, suggesting that individual toxins may act on distinct cellular targets and contribute in different ways to the complex pathophysiology of envenoming. At the same time, the high affinity and selectivity of disintegrins for integrins have made them valuable molecular tools and attractive templates for therapeutic development (Kuo et al., 2019; Mohamed Abd El-Aziz et al., 2019). However, despite their abundance at the transcript level, *E. coloratus* disintegrins have not, to the best of our knowledge, been isolated and characterised individually at the protein level. Isolation of native disintegrins in the purity and quantity required for detailed biochemical and functional analysis remains challenging because of their low abundance, structural similarity, and the complexity of the venom proteome. Recombinant expression overcomes these limitations by enabling the production of individual disintegrins in homogeneous form and sufficient quantities to unlock their structural and functional properties. This approach allows the biological properties of each toxin to be examined independently, facilitates direct comparisons between closely related variants, and provides insight into how sequence diversification within venom disintegrins translates into distinct integrin-binding and cellular activities.

Disintegrins require correct disulfide bond formation for proper folding and function. While *E. coli* is a common, cost-effective expression system, its reducing cytoplasm often leads to misfolded proteins (Baneyx, 1999). To address this, we co-expressed disintegrins with the folding catalysts Erv1p and PDI, which help the formation and isomerisation of disulfide bonds (Sohail et al., 2020). In addition, the disintegrin expression construct included an N-terminal DsbC, a bacterial disulfide isomerase that further promotes the isomerisation of mis-paired disulfide bonds, enhancing proper folding and functionality (Zhang et al., 2006). The final yields obtained in this study are comparable with other disintegrins expressed in *E. coli* (David et al., 2018), and are particularly similar to the yield reported for r-ocellatusin (0.5-1 mg/L) produced in *E. coli* BL21 cells (Sanz-Soler et al., 2012).

The anti-platelet potential of EcDis_RGD, EoDis_RGD and EcDis_KGD is similar to other disintegrins isolated from *Echis* venom. Reported IC_50_ values for ADP-induced platelet aggregation of disintegrins from the genus *Echis* can range from 63 to 360 nM including echistatin α (108 nM), echistatin β (63 nM), echistatin γ (276 nM), pyramidin A (160 nM), pyramidin B (233 nM), ocellatin (104 nM), leucogastin A (360 nM), leucogastin B (170 nM), multisquamatin (93 nM), and ocellatusin (168 nM) (Okuda et al., 2001; Smith et al., 2002). These findings suggest that the recombinant disintegrins expressed in this study adopt native-like conformations with IC_50_ values of 144 nM, 186 nM and 411 nM for EcDis_RGD, EoDis_RGD and EcDis_KGD, respectively (Table 2). Notably, EcDis_RGD and EoDis_RGD exhibited greater potency than EcDis_KGD (Figure 3). Although the KGD motif is known to be a specific binder to integrin αIIbβ3, higher IC_50_ values for ADP-induced platelet aggregation have been reported previously for KGD-bearing disintegrins. For example, ussuristatin 1 (US-1), which contains an RGD motif, inhibited ADP-, collagen-, thrombin-, and epinephrine-induced platelet aggregation with IC_50_ values ranging from 17-33 nM. In contrast, ussuristatin 2 (US-2), which contains a KGD motif, also inhibited platelet aggregation but exhibited IC_50_ values approximately ten-fold higher (Oshikawa & Terada, 1999). Barbourin, another KGD-containing disintegrin, also displayed lower potency, with an IC_50_ of 309 nM (Scarborough et al., 1991), in the same range as EcDis_KGD (411 nM). EcDis_VGD showed no detectable effect on platelet aggregation at any of the tested concentrations (500, 250, and 50 nM) in assays induced by either 2-MeS-ADP, CRP-XL, TRAP-6 (Figure 2B). This outcome was expected, as the VGD motif binds to integrin α5β1, a fibronectin receptor involved in platelet adhesion and spreading, but not in aggregation (Calvete et al., 2003; Johansson, 1997).

The higher IC_50_ values observed for CRP-XL- and TRAP-6-induced platelet aggregation compared with 2-MeS-ADP may reflect differences in the magnitude of platelet activation elicited by these agonists. Platelet aggregation is primarily mediated by activation of integrin αIIbβ3 and subsequent fibrinogen-dependent platelet–platelet bridging; therefore, inhibition of αIIbβ3 by disintegrins can effectively prevent the fibrinogen-mediated aggregation response. However, CRP-XL and TRAP-6 are relatively strong agonists that induce extensive platelet activation, including rapid activation of a large pool of integrin αIIbβ3, pronounced release of amplification factors such as ADP and TxA_2_, integrin αIIbβ3 receptor clustering and outside-in signalling. In contrast, 2-MeS-ADP is a considerably weaker agonist and induces more limited platelet activation (Zou et al., 2024). Consequently, the aggregation response by CRP-XL and TRAP-6 involves a more complex and extensive activation state than that elicited by 2-MeS-ADP and may explain why complete inhibition of CRP-XL- and TRAP-6-induced aggregation was not achieved even at maximal disintegrin concentrations.

Our data confirm a disintegrin- and concentration-dependent modulation of platelet aggregation, highlighting distinct functional properties among the recombinant proteins. Taken together, these comparisons support the validity of EcDis_RGD, EoDis_RGD and EcDis_KGD as biologically active integrin αIIbβ3 inhibitors. While platelet aggregation assays provide essential insights into the antiplatelet potential of disintegrins, they offer only a partial view of their biological activity, as disintegrins can interact with a wide range of integrin receptors beyond integrin αIIbβ3. Hence, a cell migration assay was performed to help elucidate interactions with non-platelet integrins. The scratch wound healing assay (Figure 4) revealed marked differences among the disintegrins in their ability to modulate endothelial cell migration. EcDis_RGD consistently inhibited HUVEC migration, whereas EoDis_RGD produced only modest inhibition at the highest concentration tested. EcDis_KGD and EcDis_VGD showed no detectable activity. These results show that RGD-containing disintegrins may interfere more effectively with integrin receptors involved in endothelial cell motility, such as integrin αvβ3, αvβ5, and α5β1, which are also implicated in tumour growth and angiogenesis (David et al., 2018). The differences observed between EcDis_RGD and EoDis_RGD indicate that biological activity is determined not only by the RGD integrin-binding motif itself but also by the structural context in which it is presented. Previous studies have shown that integrin affinity and specificity are strongly influenced by the conformation of the integrin-binding loop and the surrounding amino acid residues, allowing disintegrins with identical motifs to display distinct biological activities (Calvete et al., 2005; Kim et al., 2005). Together, these findings highlight the potential of RGD-containing disintegrins as modulators of integrin-dependent cellular processes and support further investigation of their anti-migratory and anti-angiogenic properties for biomedical applications.

The specificity of the KGD motif for integrin αIIbβ3 could explain the inability of EcDis_KGD to inhibit HUVEC migration, as this integrin is predominantly expressed on platelets and is not a mediator of endothelial cell motility (Huang et al., 2019). At first glance, the lack of a significant effect of EcDis_VGD on HUVEC migration appears surprising, as previous studies have reported that the VGD motif binds to integrin α5β1 (Calvete et al., 2003) - a fibronectin receptor involved in cell migration and adhesion (Johansson, 1997). However, all VGD-containing disintegrins described to date are heterodimeric molecules, and the VGD-containing subunit appears to contribute only modestly to integrin binding (Almeida et al., 2023; Calvete et al., 2003). For example, the VGD-containing subunit EC3A, after reduction and ethylpyridylethylation, inhibited adhesion of K562 cells (a human haematopoietic cell line) to fibronectin with an IC_50_ value of approximately 30 μM, while EC3 (the heterodimeric form with VGD and MLD-containing subunits) presented an IC_50_ of 150 nM (Marcinkiewicz et al., 1999). Therefore, the absence of a detectable effect of EcDis_VGD at concentrations up to 1 μM (Figure 4) may reflect its comparatively low affinity for integrin α5β1 receptors and the possibility that higher concentrations are required to disrupt integrin-mediated cell migration.

While the biological activities of these *Echis* disintegrins are broadly consistent with previously characterised disintegrins, our study provides a useful comparative framework for investigating the determinants of integrin specificity and selectivity. Recombinant production enabled individual disintegrins with different integrin-binding motifs to be evaluated under comparable experimental conditions, revealing differences in their ability to inhibit platelet aggregation and endothelial cell migration. The distinct functional profiles observed for the RGD-, KGD-, and VGD-containing disintegrins, despite their moderate-to-high sequence identity, suggest that receptor recognition is influenced not only by the canonical binding motif but also by its surrounding sequence and structural context. Such insights are relevant to drug design, as understanding the molecular features that favour interaction with specific integrins could enable the rational modification of disintegrin scaffolds to improve receptor selectivity and therapeutic properties (Kuo et al., 2019), similar to the development paths of eptifibatide and tirofiban (Phillips & Scarborough, 1997; Lynch et al., 1995). By enabling naturally occurring disintegrin variants to be compared under identical experimental conditions, recombinant production provides a foundation for dissecting how sequence variation surrounding canonical integrin-binding motifs influences integrin recognition and biological activity. These proteins therefore expand the molecular diversity available for defining determinants of selective integrin recognition and for engineering integrin-targeting molecules. Ocellatusin can theoretically be generated either by proteolytic processing of a P-II SVMP precursor (EoSVMP_PII) or by direct translation of a short-coding disintegrin transcript (same sequence as EoDis_RGD), and although it has been suggested that the short-coding message represents the major precursor of ocellatusin (Wagstaff et al., 2009), the relative contribution of these two biosynthetic pathways in *Echis ocellatus* remains unknown. Both mechanisms have been described in viperid snakes (Fox & Serrano, 2008; Juárez et al., 2006) and current transcriptomic and proteomic data are insufficient to determine the predominant source of the mature toxin.

While native ocellatusin and the recombinant disintegrin EoDis_RGD exhibited comparable inhibitory potency, its recombinant SVMP PII precursor showed a weaker inhibitory profile, even following autoactivation of the metalloprotease domain and release of the disintegrin (Figure 5C). Although alternative processing products cannot be completely excluded, MALDI analysis suggests that the disintegrin species generated from the PII SVMP has the same N-terminus as ocellatusin and retains an additional 29 amino acid residues corresponding to the C-terminal region of the disintegrin domain and the Avi-tag (Figure S6). However, MS analysis is not quantitative, and the Coomassie-stained reducing SDS-PAGE indicates that autoactivation is incomplete, suggesting that the preparation likely contains a mixture of processed and unprocessed protein. The lower apparent inhibitory activity of the autoactivated PII-SVMP preparation is therefore consistent with incomplete generation of a fully active disintegrin species. However, the heterogeneous nature of the processed sample precludes conclusions regarding the intrinsic activity of the SVMP-derived disintegrin. Further studies employing purified processing intermediates and mature disintegrins will be required to determine whether the additional C-terminal residues directly influence integrin recognition or inhibitory potency.

## 5. Conclusion

In this study, *Echis* disintegrins were successfully expressed in *E. coli* and purified, enabling their biochemical and functional characterisation. The recombinant disintegrins displayed biological activities consistent with their predicted integrin-binding specificities. RGD-containing disintegrins (EcDis_RGD, followed by EoDis_RGD) were the most potent inhibitors of platelet aggregation and endothelial cell migration. The KGD-containing disintegrin selectively inhibited platelet aggregation but not wound healing. EcDis_VGD did not inhibit platelet aggregation, and, unexpectedly, also showed no detectable effect on endothelial migration under the conditions tested. Our findings also highlighted the importance of both the integrin-binding motif and its surrounding sequence context in determining disintegrin function. Furthermore, comparison of native ocellatusin with its recombinant counterparts demonstrated that the recombinant disintegrin faithfully reproduces the biological activity of the native toxin, validating the recombinant expression strategy as a robust platform for the production of disintegrins with native-like activity. In contrast, the lower apparent inhibitory activity of the autoactivated PII-SVMP preparation is consistent with incomplete generation of a fully active disintegrin species. Collectively, these findings improve our understanding of disintegrin structure-function relationships and support the use of these recombinant venom toxins as valuable tools for studying integrin biology and for the development of therapeutic applications.

## Data Availability Statement

All data generated or analysed during this study are included in the manuscript and supporting files.

## Acknowledgment

The authors thank all members of the Schaffitzel, Berger, Poole, Hers and Casewell teams for discussions and support. The Wolfson Bioimaging Facility team is acknowledged for support with the Incucyte system, and the Liverpool School of Tropical Medicine Herpetarium team for the provision of venom samples used for toxin isolation.

## Authors contributions

Conceptualisation: I.A.C., C.S., A.W.P.; Methodology: I.A.C., S.H., I.H., R.V., A.W.P., C.S.; Validation: I.A.C., I.H.; Formal Analysis: I.A.C., T.C., L.C., A.W.P., L.Q, N.R.C., I.B., C.S.; Investigation: I.A.C., S.H., T.C., B.R., D.S., M.W., R.S., A.R., C.W., D.W.; Supervision: C.S. A.W. P.; Writing – Original Draft Preparation: I.A.C, C.S.; Writing – Review & Editing: C.S., A.W.P., I.H., I.B., N.R.C., L.Q.

---

**Figure S1.**
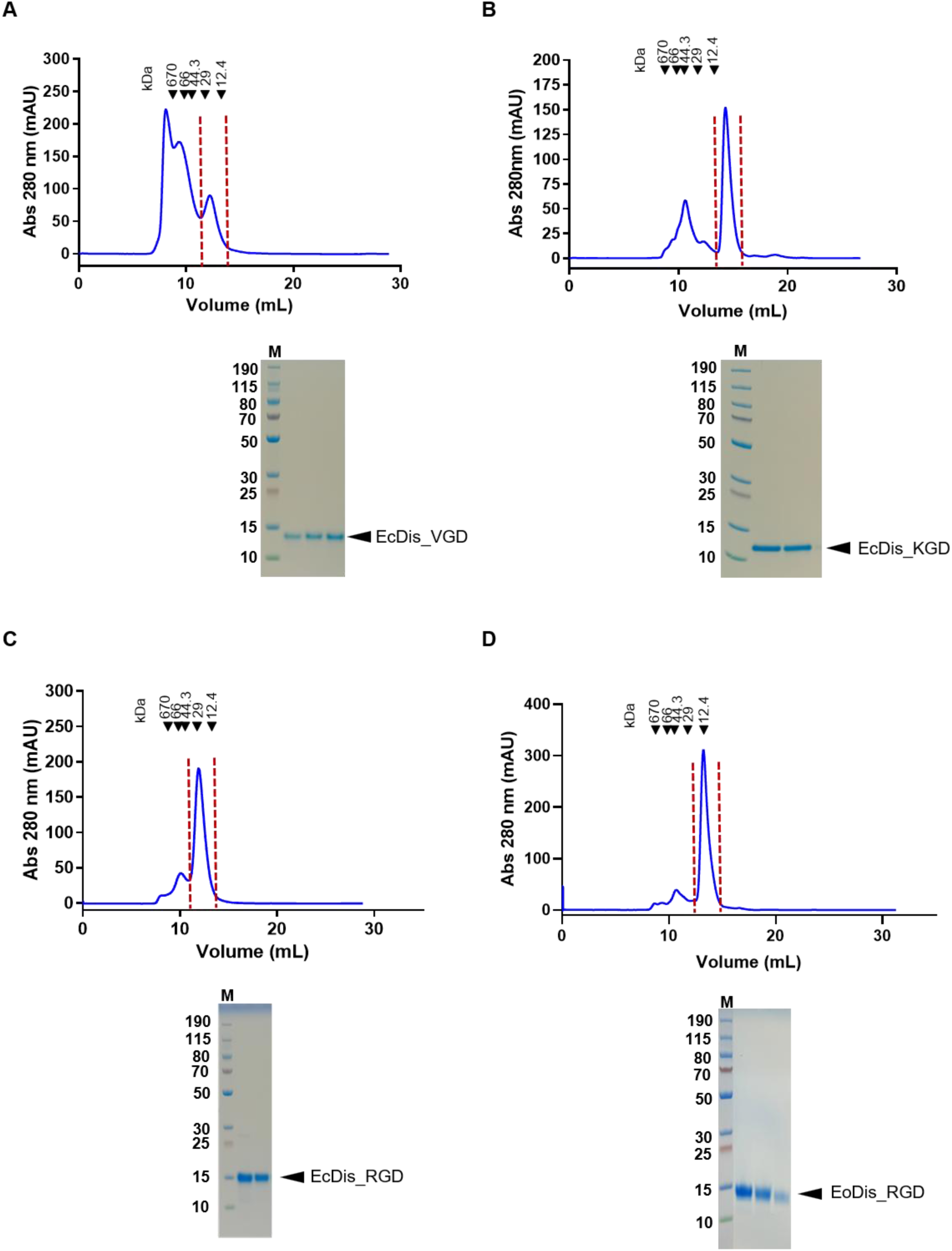
SEC chromatograms of disintegrin purification. Size exclusion chromatogram of **A)** EcDis_VGD, **B)** EcDis_KGD, **C)** EcDis_RGD and **D)** EocDis_RGD, with the corresponding disintegrin peak highlighted by dashed red lines and Coomassie-stained SDS-PAGE gel (reducing conditions). M: PageRuler™ Plus Prestained Protein Ladder.

**Figure S2.**
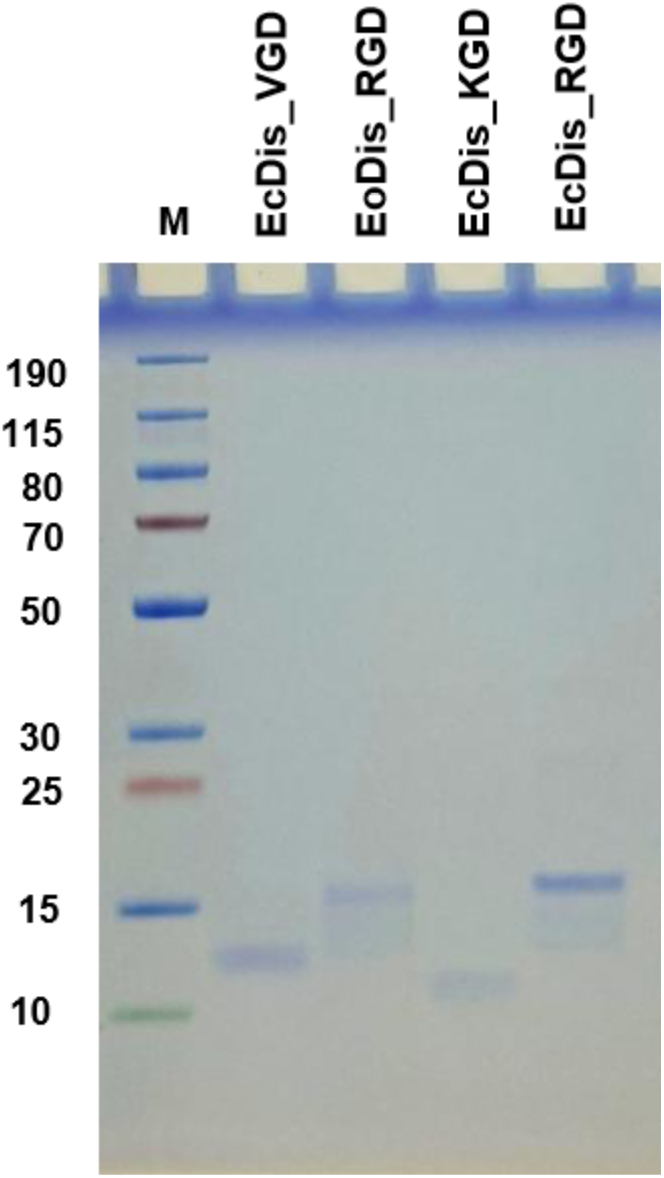
Non-reducing Coomassie-stained SDS-PAGE analysis of purified EcDis_VGD, EoDis_RGD, EcDis_KGD, and EcDis_RGD, following size-exclusion chromatography. M: PageRuler™ Plus Prestained Protein Ladder.

**Figure S3.**
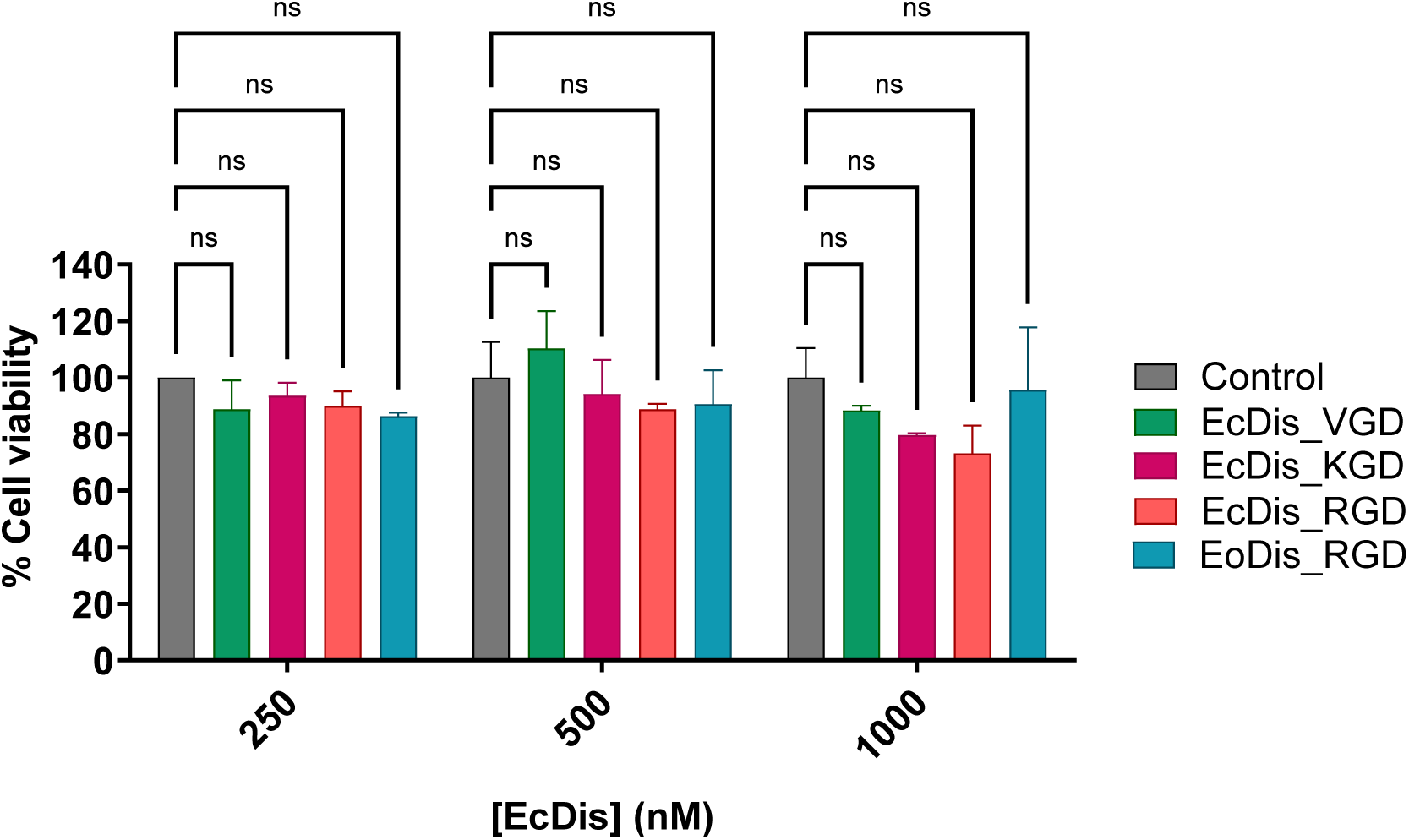
HUVEC cell viability. MTT cytotoxicity assay showing HUVEC cell viability compared to control after 23 h incubation with EcDis_VGD (green), EcDis_KGD (magenta), EcDis_RGD (red) and EoDis_RGD (dark teal). Control is shown in dark grey. Bars show the results of two independent repeats. Statistical analysis was obtained using Two-way ANOVA and Dunnett’s Multiple Comparison post-test, in GraphPad Prism 11.

**Figure S4.**
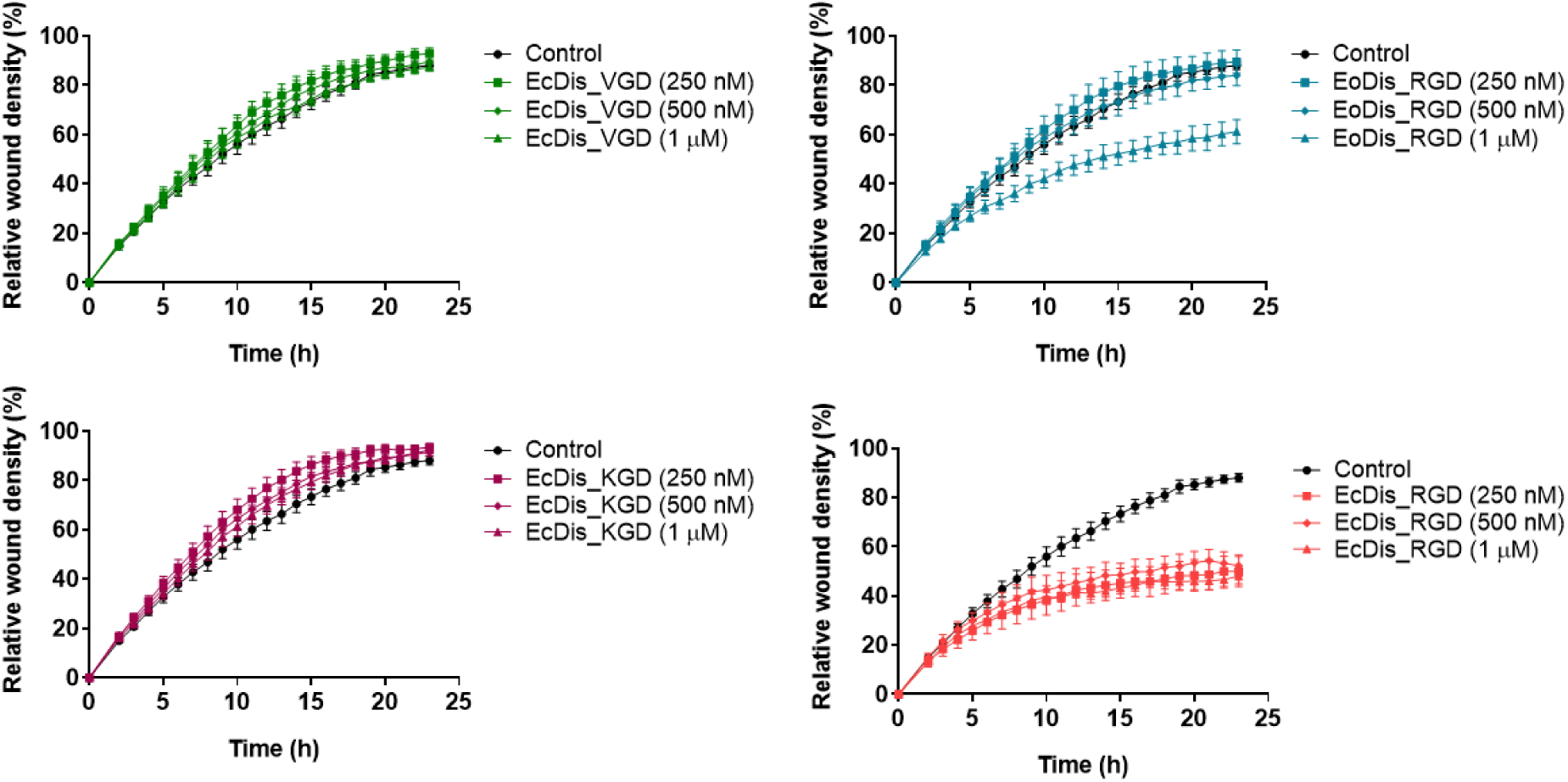
Relative wound density (%) versus time (h). Disintegrin-treated cells were tested at 250 nM (squares), 500 nM (diamonds), and 1 µM (triangles). Control curves (without disintegrins) are shown in black. EcDis_VGD is shown in green, EoDis_RGD is shown in dark teal, EcDis_KGD in magenta and EcDis_RGD in red.

**Figure S5.**
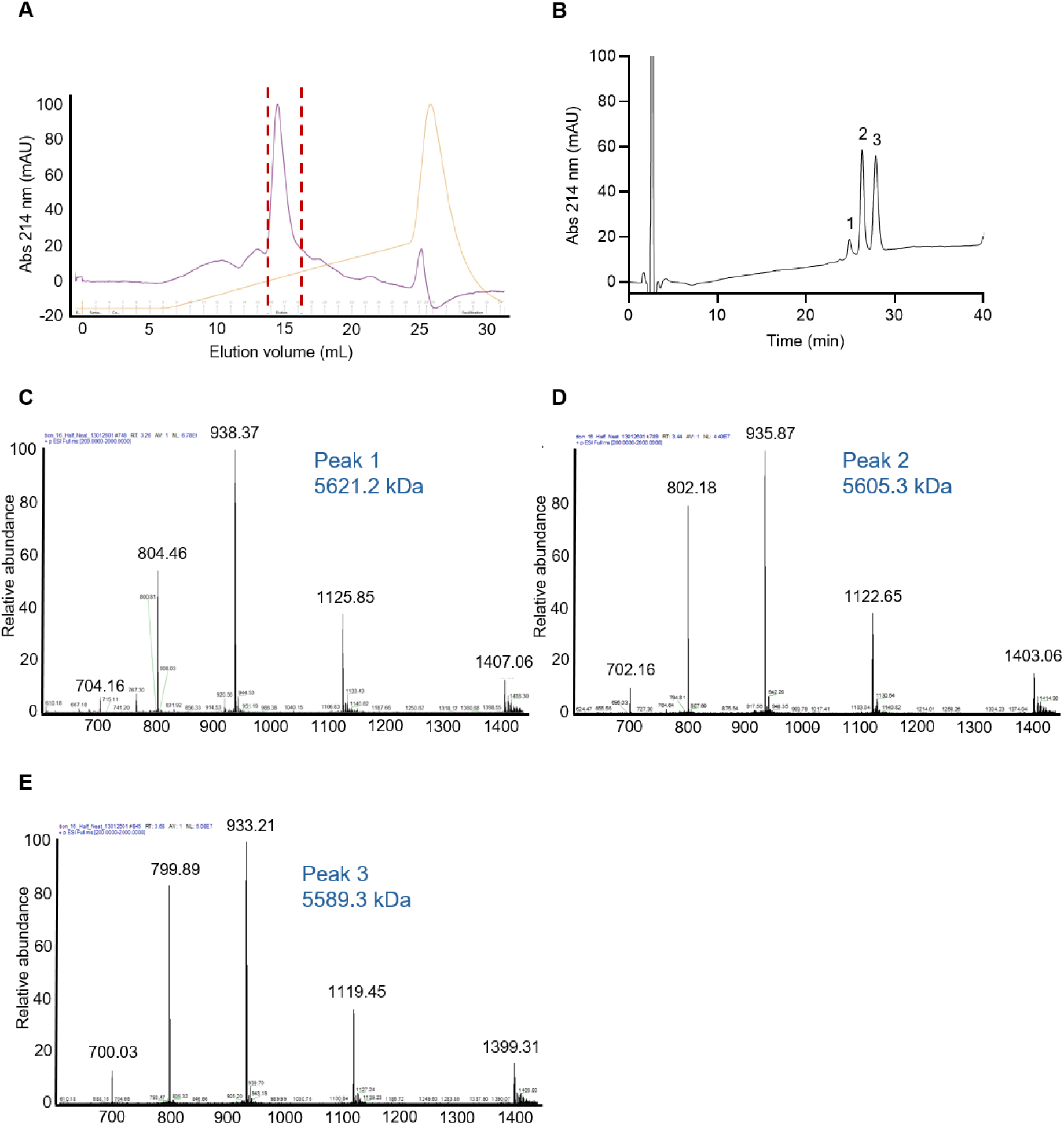
Purification and identity confirmation of ocellatusin. **A)** AIEX profile obtained during ocellatusin purification. **B)** RP-HPLC profile of the major peak obtained from AIEX chromatography. Peaks 1–3 were analysed by mass spectrometry. Intact mass spectra of **C)** Peak 1, **D)** Peak 2, and **E)** Peak 3 corresponded to ocellatusin containing two oxidised methionine residues, one oxidised methionine residue, and unmodified ocellatusin, respectively. Intact mass spectra were acquired by electrospray ionisation (ESI) in positive-ion mode using an Orbitrap mass spectrometer, with full-scan spectra collected over m/z 200– 2,000 at a resolution of 120,000 (at m/z 200).

**Figure S6.**
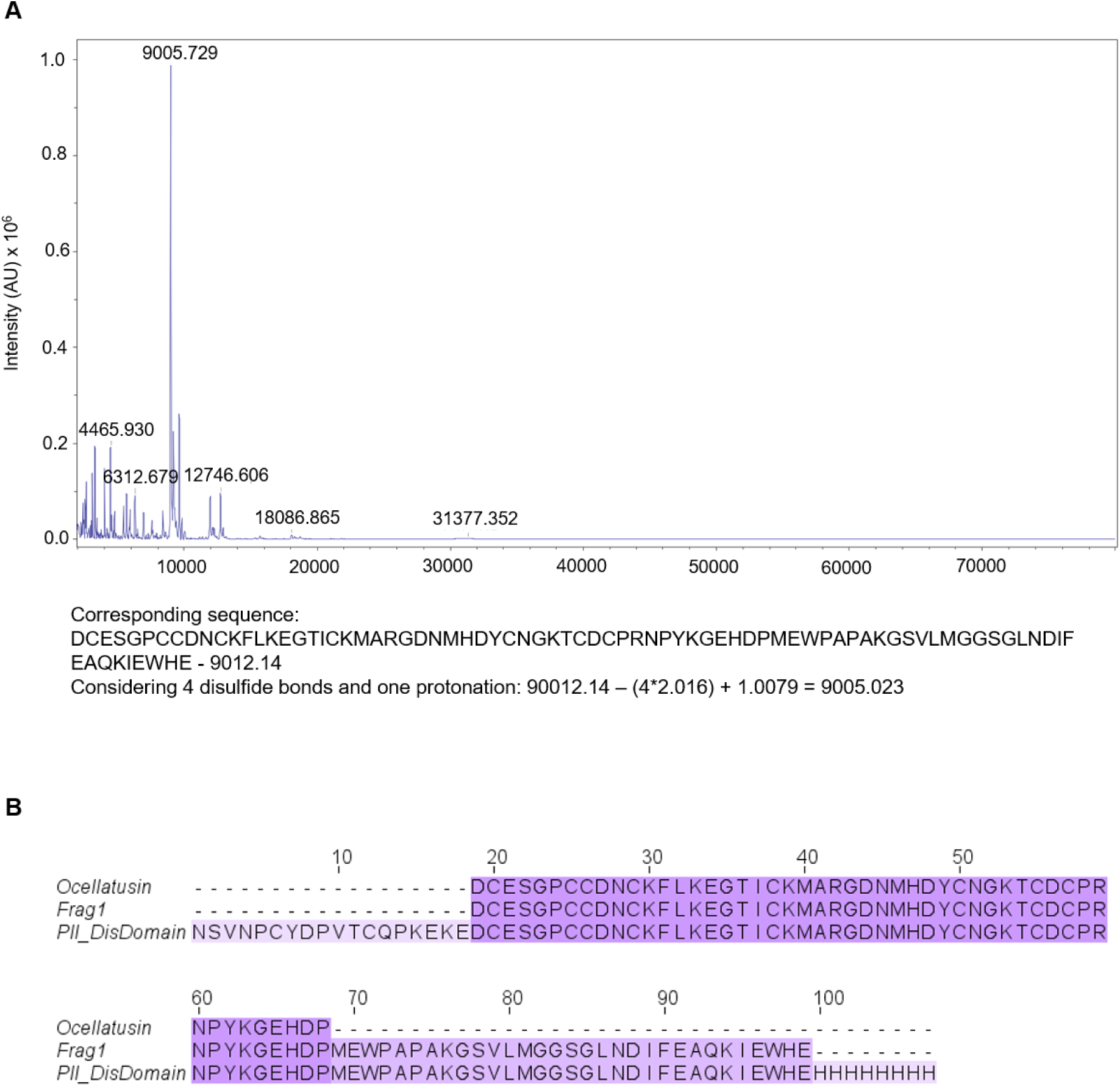
Intact mass analysis and sequence comparisons of EoSVMP_PII_zym_ autoactivation product. **A)** MALDI-TOF MS analysis of the EoSVMP_PII_zym_ autoactivation reaction, showing the protein and peptide components present in the sample. Mass spectra were acquired in positive linear mode over a mass range of 2–80 kDa. The sequence identified from the sample is shown below the corresponding mass spectrum. **B)** Sequence alignment of native ocellatusin (UniProt ID Q3BER1), the disintegrin fragment (Frag1) identified following EoSVMP_PII_zym_ autoactivation, and the disintegrin domain of EoSVMP_PII (PII_DisDomain, UniProt ID Q14FJ4). Multiple sequence alignment was generated by Clustal Omega and visualised using Jalview.

